# A set of Columbia-0-specific single nucleotide polymorphism markers for the genetic analysis of natural variation in *Arabidopsis thaliana*

**DOI:** 10.1101/153197

**Authors:** Ulrich Lutz, Claus Schwechheimer

**Author notes:** Corresponding author: Claus Schwechheimer.

## Abstract

Genetic markers are important tools for the study of natural and induced genetic variation. Due to the strong increase in the number of genome sequences, an overview of the genomic diversity of many natural strains from individual species could be gained. This allows for the design of markers for flexible use and cost-efficient small scale genetic studies requiring minimal laboratory and bioinformatics effort. Here, we describe 140 single nucleotide polymorphism (SNP) markers with genome-wide distribution that discriminate between the genotype of the common *Arabidopsis thaliana* reference accession Columbia-0 (Col-0) and the majority of *Arabidopsis thaliana* accessions that have been sequenced to date. We designed, generated and validated the markers using the kompetitive allele-specific PCR (KASP) technology and made all 140SNPvCol marker assays publicly available through a service provider. Through the integration of available genomic SNP allele information of 1,135 accessions, we found that 120 of these 140 markers could detect non-reference alleles in 647 accessions and more than 100 markers showed non-reference alleles in 1,094 accessions. We further show that the marker set can be used for the verification or identification of accessions of unknown identity. As the KASP methodology is very flexible, an optimal marker subset can be easily selected among the available 140SNPvCol markers presented here to analyze genetic combinations of Col-0 with any other accession.

## Introduction

Forward genetics techniques serve to identify the underlying genetic loci contributing to phenotypic variation. Genetic analyses benefit from the presence of intraspecific sequence polymorphisms such as single nucleotide polymorphisms (SNPs) or small nucleotide insertions or deletions (Indels) that can be used for distinguishing genomic regions and genetic mapping (Lukowitz et al., 2000; Alonso-Blanco et al., 2009). In the model plant *Arabidopsis thaliana* (Arabidopsis), many positional cloning strategies have been described to date and genetic marker sets have been designed to provide high mapping resolution and to discriminate between the most commonly used accessions such as Columbia-0 (Col-0), Cape verde island-0 (Cvi-0), Landsberg *erecta* (L*er*), and Wassilewskaya-0 (Ws-0) (Alonso-Blanco et al., 2009) (Peters et al., 2003; Pacurar et al., 2012).

In the past decade, an increasing number of studies have been dedicated to the understanding of natural variation in Arabidopsis (Weigel and Nordborg, 2005; Weigel, 2012; The 1001 Genomes Consortium, 2016). As opposed to forward genetic approaches, where phenotypic diversity is being studied following mutagenesis of a reference species, natural variation studies are concerned with the genetic causes underlying the phenotypic diversity within a species as a result of evolutionary adaptations to specific environments. The analysis of naturally occurring variants challenges genetic analyses since it requires tools that allow discriminating between the polymorphisms present between essentially any pair of genomes.

Since SNPs are very frequent, even between genomes of very related individuals from one species, they are able to provide the highest possible resolution when the best discrimination between two related genomes is desired. In Arabidopsis, coordinated next generation sequencing- (NGS-)based sequencing efforts have recently provided the genome sequences of 1,135 Arabidopsis accessions and thereby, an overview of the pangenomic diversity (The 1001 Genomes Consortium, 2016). In Arabidopsis, the accessions sequence to date has a median of 439,145 SNPs when compared to the genome of the reference accession Col-0. The current version of the Arabidopsis pangenome detects one SNP every eleven bp and this renders Arabidopsis the organism with the densest variant map of any organism analysed to date (The 1001 Genomes Consortium, 2016).

Various SNP genotyping technologies allow for the genome-wide determination of SNP alleles in a single experiment (Konieczny and Ausubel, 1993; Tabone et al., 2009). In Arabidopsis, different SNP detection technologies have been established such as the Sequenom Mass Array (compact) system for the parallel genotyping of 149 SNPs (Platt et al., 2010), the Golden Gate Genotyping Assay for 384 SNPs (Simon et al., 2012; Agrena et al., 2013), the SNPlex technique for sets between 96 and 320 SNPs (Simon et al., 2008) (Pico et al., 2008; Gomaa et al., 2011; Mendez-Vigo et al., 2013), a matrix-assisted laser desorption/ionization time-of-flight (MALDI-ToF) analysis for 115 SNPs (Torjek et al., 2003; Schmid et al., 2006) and DNA-array systems that can genotype tens of thousands of SNPs (Gresham et al., 2010; Yuan et al., 2016).

On the other side of the spectrum, NGS-based genotyping-by-sequencing approaches make it possible to potentially detect all SNPs of segregating genomes of mapping populations in a multiplexed sequencing effort, at least for small genomes (James et al., 2013; Rowan et al., 2015). However, these pipelines are not applicable to small-scale genetic studies as sample and library preparation is complex and costly compared to other methods. Further, genotyping-by-sequencing approaches require substantial laboratory and bioinformatics knowledge or support that may not be present in individual laboratories.

Kompetitive allele-specific PCR (KASP) is a comparatively new technique for SNP genotyping with demonstrated usefulness in Arabidopsis (Wijnker et al., 2012; Semagn et al., 2014; Thomson, 2014). KASP is based on allele-specific oligo extension combined with a fluorescence resonance energy transfer (FRET) system for signal detection. Compared to existing small-scale SNP genotyping platforms, the KASP technology provides high flexibility, while being cost-efficient, with a fast turnover and high accuracy. KASP only requires a thermocycler for real-time quantitative PCR measurements, which is available in most modern molecular biology laboratories.

Here, we introduce the KASP technology-based genome-wide 140SNPvCol marker set that specifically allows discriminating between the genome of the widely used reference accession Col-0 and that of the majority of the 1,135 Arabidopsis accessions released from the 1001 Genomes project (The 1001 Genomes Consortium, 2016). We made this marker set available to the public through a service provider, restricting the technical requirements to DNA preparation before the analysis and allowing for minimal bioinformatics requirements after the analysis.

## Results

### Selection of 140 SNP markers for the 140SNPvCol marker set

With the aim of generating a marker set for genetic analyses that distinguishes the reference accession Col-0 from Kil-0, an accession that had become relevant in one of our previous studies (Lutz et al., 2015), we selected SNPs between Col-0 and Kil-0 from the TAIR10 assembly for Col-0. We restricted our selection to SNPs with a low frequency of the Col-0 allele by investigating allele frequency using the genome sequence information from a randomly chosen small subset of accessions (The 1001 Genomes Consortium, 2016). In this way, we obtained a set of 140 bi-allelic SNPs designated 140SNPvCol (Fig. 1). These SNP markers were equally distributed over the genome and we expected them to be highly informative not only for comparisons between Col-0 and Kil-0 but also for comparisons to a large proportion of the sequenced Arabidopsis accessions (Fig. 1). In 140SNPvCol, the average marker distance between two neighbouring markers was 871 kb and ranged between 101 kb (between CK_SNP_36 and CK_SNP_33 on Chromosome 1) and 5,478 kb (between CK_SNP_15 and CK_SNP_18 on Chromosome 1) (Fig. 1, Suppl. Table 2).

**Figure 1:**
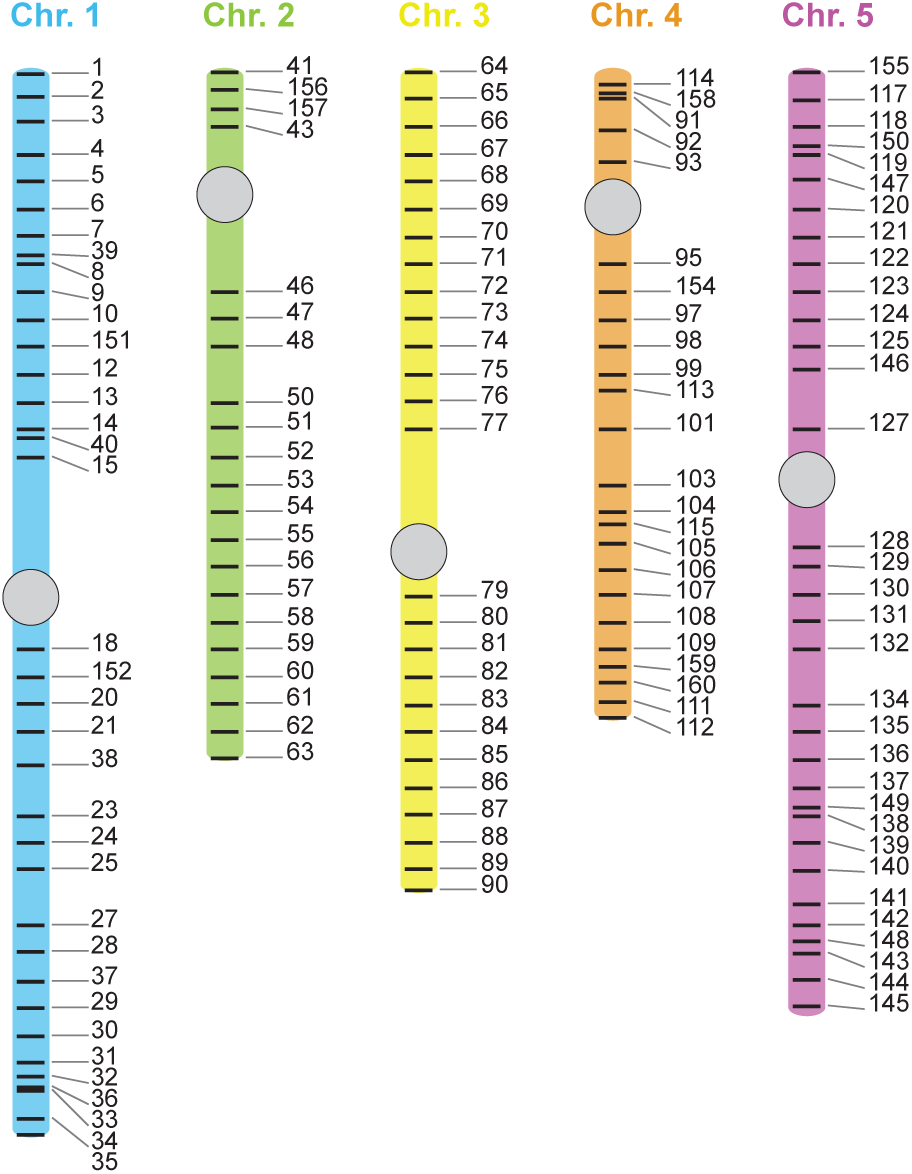
Physical position of the 140SNPvCol markers on the Col-0 reference genome. 140SNPvCol marker IDs (only ID shown) along the five Arabidopsis chromosomes. Centromeres are indicated with a grey circle.

### Validation of the 140SNPvCol markers

To make the 140SNPvCol SNP marker set amenable to routine genetic analyses for a larger group of users, we established KASP-based SNP assays (Semagn et al., 2014; Thomson, 2014). To validate the KASP-based assays for the entire 140SNPvCol marker set, we genotyped the accessions Col-0 (1001 Genomes project accession ID6909) and Kil-0 (ID7192), allowing us to unambiguously determine all genotypes for the 140 markers. They segregated between Col-0 and Kil-0 and were homozygous, as expected for inbred wild type strains (Suppl. Table 3). We also genotyped L*er*-0 (ID7213), another commonly used reference accession for mutant analyses, and were able to unambiguously determine all genotypes with the 140SNPvCol markers. When we compared the L*er*-0 140SNPvCol genotypes with the respective L*er*-0 SNP genotypes deposited in the 1001 Genomes project database (http://1001genomes.org/), we noted the absence of one genotype (ambiguous allele N) but the presence of 139 unambiguously determined genotypes that all coincided with the genotypes determined by the 140SNPvCol markers (Suppl. Tables 3 and 4). We concluded that the designed 140SNPvCol marker set can be used for the efficient genotyping of Arabidopsis accessions.

### 140SNPvCol marker allele frequency among 1,135 *Arabidopsis thaliana* accessions

We next determined whether the 140SNPvCol marker set was useful for the analysis of a larger set of Arabidopsis accessions. To this end, we determined the allele frequencies for the 140SNPvCol markers from the available Arabidopsis genome sequences. Among the 1,135 sequenced accessions, we found that 136 of the 140SNPvCol markers showed minor frequencies (< 50%) of the Col-0 reference (R) allele and major frequencies (≥ 50%) of the non-reference (NR) allele. Only four markers (CK_SNP_158, 95, 106, 128) showed a higher frequency of the reference R allele (Fig. 2). Among the 158,900 different genotypes of the 140 markers among the 1,135 accessions, only 11.5% (18,265) of genotypes corresponded to the R allele but 85% (134,378) corresponded to the NR allele. Only 6,257 (3.9%) were not determined and were therefore, designated as ambiguous alleles (N). The CK_SNP_156 showed an especially high number of ambiguous genotypes in the available SNP data, maybe due to technical difficulties in read mapping at this specific locus (Fig. 2 and Suppl. Table 4). Since 85% of the selected markers were divergent between the reference accession Col-0 and the analysed 1,135 accessions, we concluded that the 140SNPvCol marker set was highly informative to discriminate between Col-0 and a large number of accessions.

**Figure 2:**
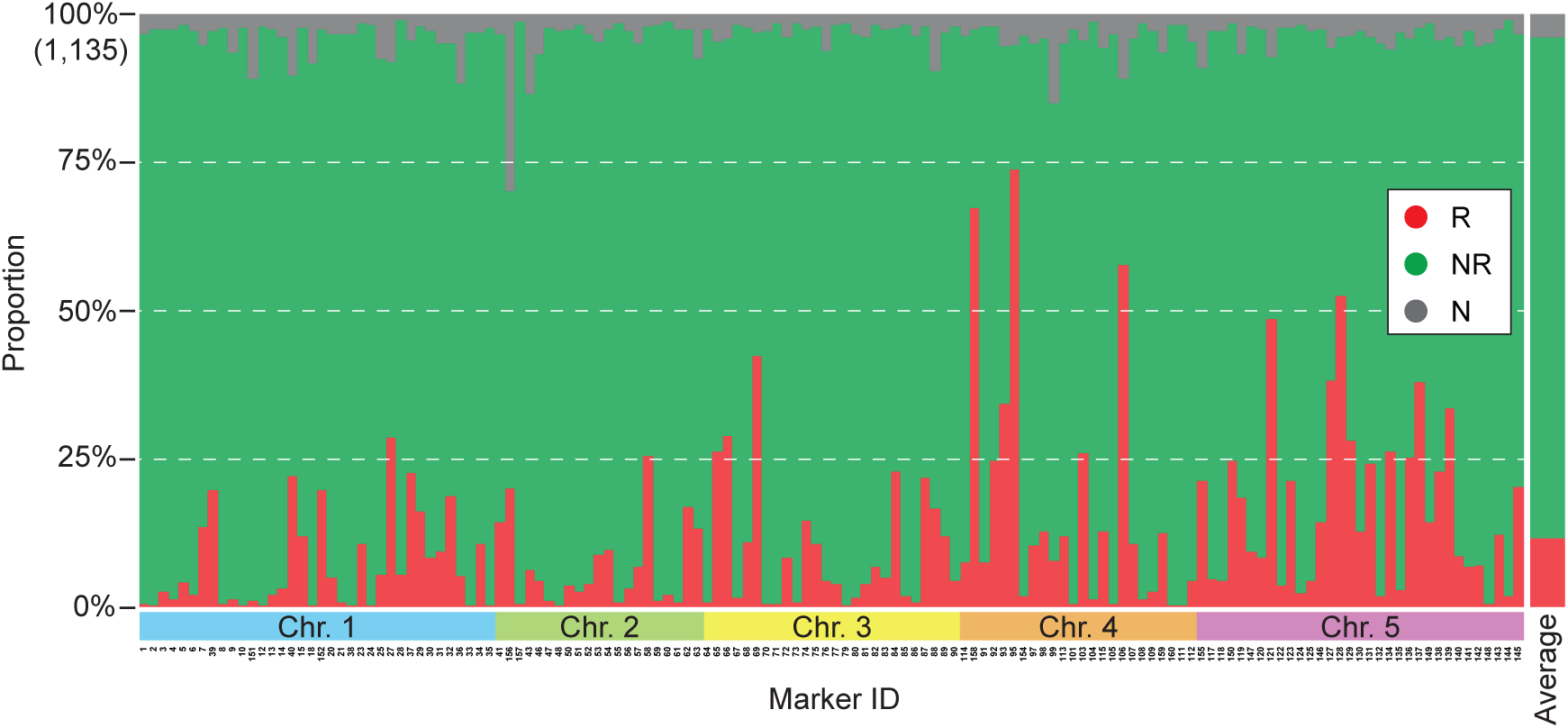
Allele frequencies of the 140SNPvCol markers among 1,135 accessions. Graph showing the proportion of Col-0 reference (R) alleles (red), non-reference (NR) alleles (green) and ambiguous (N) alleles (grey). Proportions are shown with grey horizontal dotted lines, with 100% representing all 1,135 accessions. The average of the number of R, NR, and N alleles among all 140SNPvCol markers and 1,135 accessions is shown on the right.

We next analysed the allelic composition of each of the 1,135 accessions. The 140SNPvCol marker set was originally designed to be maximally informative to discriminate between Col-0 and Kil-0. Therefore, Col-0 (ID6909) and Kil-0 (two samples deposited: ID7192 and ID5748) showed, as expected, the lowest and highest number of NR alleles among the 1,135 accessions, respectively (Fig. 3). Strikingly, more than 100 among the 140 SNPs showed the NR allele in 1,094 accessions (96.4%) and 120 SNPs showed the NR allele in 647 (57%) accessions. In this analysis, none of the ambiguous alleles were considered, such that the frequencies of NR alleles, as stated above, are most likely underestimations (Fig. 3). Only four of the 1,135 accessions had fewer than 60 NR alleles (TÄL07, ID6180; Lan-0, ID7208; H55, ID7461; UKSE06-252, ID5104) (Fig. 3). Also, the other commonly used reference accessions, Cvi-0 (ID6911) and Ws-0 (deposited as Ws-0.2, ID7396), showed 131 and 121 NR alleles, respectively. Therefore, the 140SNPvCol marker set is also useful for the genotyping of Cvi-0/Col-0 and Ws-0/Col-0 mapping populations. In summary, we concluded that the 140SNPvCol marker set can be used for the genetic discrimination between Col-0 and most of the 1,134 sequenced Arabidopsis accessions.

**Figure 3:**
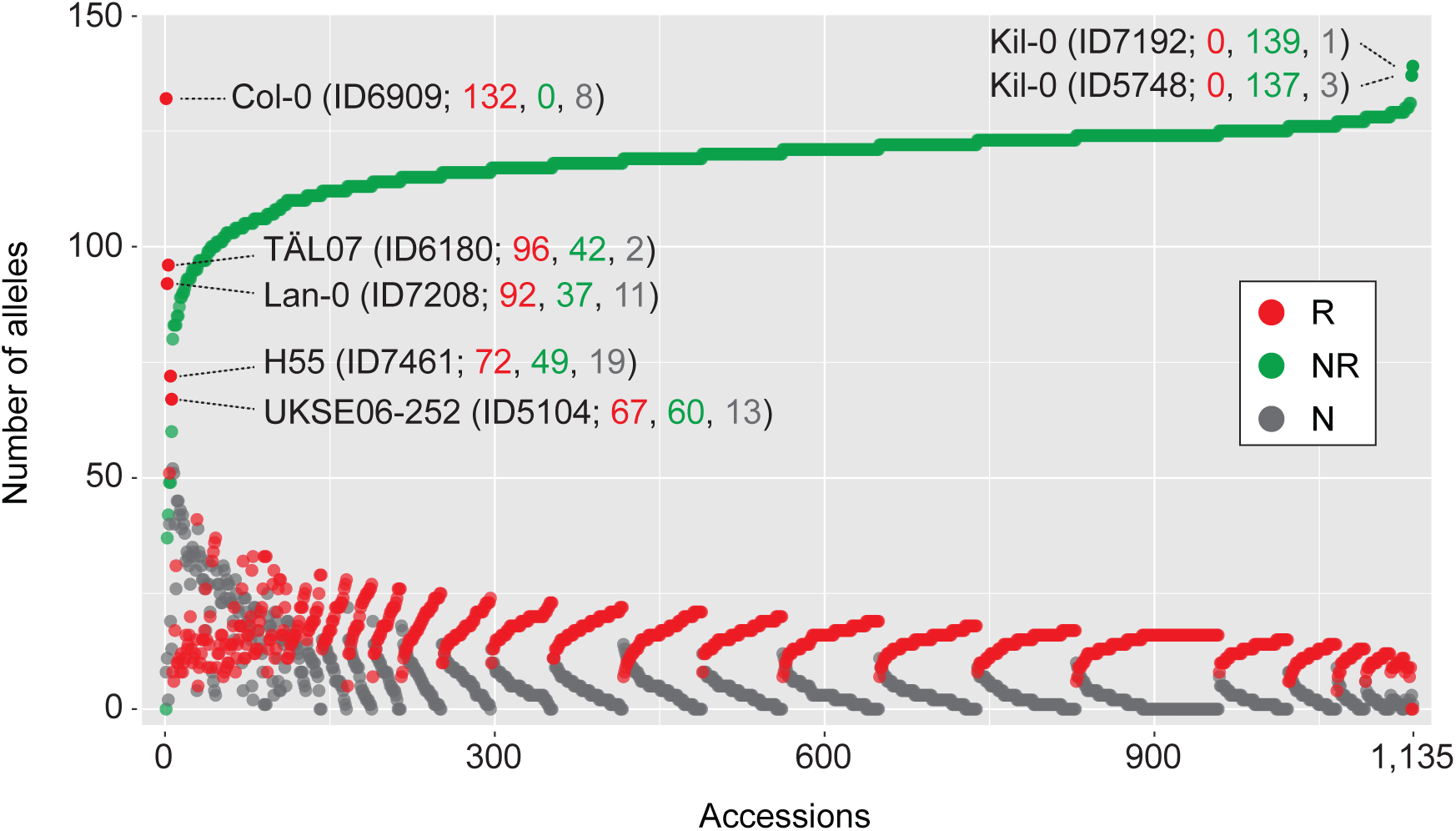
Allelic 140SNPvCol composition of each of the 1,135 accessions. Plot showing the numbers of Col-0 reference (R) alleles (red), non-reference (NR) alleles (green) and ambiguous (N) alleles (grey). The accessions were first sorted by the number of NR alleles and subsequently by the number of R alleles. The x-axis indicates a running accession number. This does not correspond to the accession ID of the 1001 Genomes project but can be retrieved from Suppl. Table 4.

### Identification of accessions through determination of the 140vC haplotype

We next asked whether the 140SNPvCol marker set could be used to distinguish between any pair of genotypes. To this end, we calculated the pairwise distances between all 643,545 possible pairs among the 1,135 accessions, while excluding all ambiguous genotypes (N). As expected, a large proportion of pairs showed a high number of shared, thus non-informative, alleles, reflecting the overall low minor allele frequency of the Col-0 allele among the 140SNPvCol SNPs (Fig. 4A). Only 1.23% (7,927) of the accession pairs showed between 0 and 70 shared alleles and a maximum of 5.67% (36,502) of pairs showed 111 shared alleles. Hence, as was expected from its Col-0-specific design, the marker set is comparatively uninformative for the discrimination between pairs among the other 1,134 accessions.

**Figure 4:**
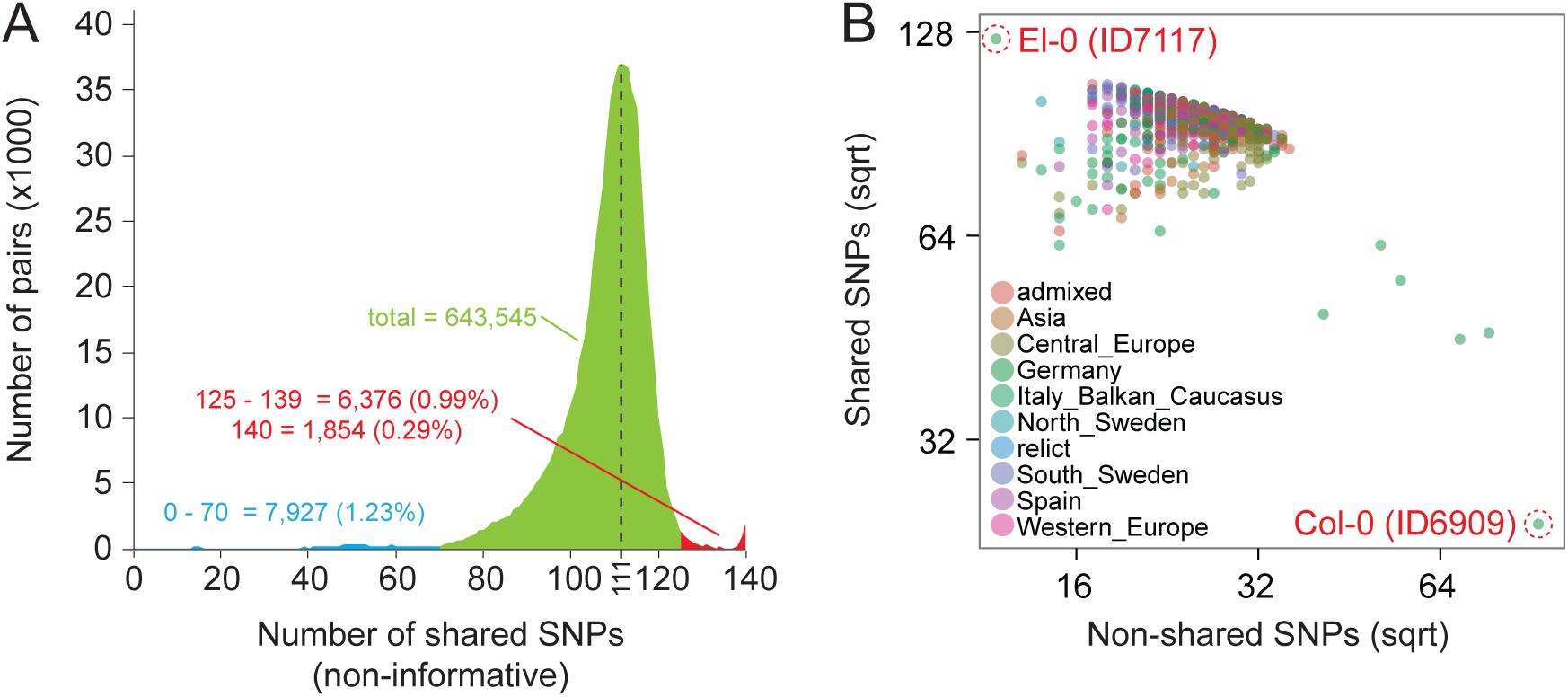
Pairwise genetic distance among 1,135 accessions. A. Graph displaying the frequency of the pairwise genetic distances between 140SNPvCol genotypes of the 1,135 sequenced Arabidopsis thaliana accessions. Fractions are shown in blue and red, the total number of all pairs in green, and the maximum number of pairs, which was found for 111 shared SNPs, is shown with a vertical dotted line. B. Comparison of genotypes of 125 markers of the unknown accession Unk (Lutz et al., 2015) with the corresponding marker alleles of 1,135 accessions. The accession with the highest number of shared SNPs (El-0) and the highest number of non-shared SNPs (Col-0) are highlighted with circles. x- and y-axis scales were square root-transformed as indicated. The colour code represents the ADMIXTURE genetic group membership of each accession as presented in (The 1001 Genomes Consortium, 2016).

We, however, also detected that only 0.99% (6,376) of the accession pairs (643,545) shared between 125 and 139 shared alleles and only 0.29% (1,854) of the accession pairs showed 140 shared alleles and, hence, had an identical 140SNPvCol haplotype (Fig. 4A and Suppl. File 2). Even though this proportion may be slightly biased because ambiguous N alleles were ignored, we concluded that the 140SNPvCol markers were useful to determine or verify the identity of accessions, or at least to assign a genetic group membership by determining the specific 140SNPvCol haplotype. To test this possibility, we genotyped an accession of unknown identity, that we designated Unknown (Unk) (Lutz et al., 2015). When Unk was genotyped, 125 of the 140SNPvCol markers provided informative readouts. When we determined the number of shared alleles among the 1,135 accessions, we found a perfect consensus in all 125 genotypes between Unk and the accession El-0 (ID7117), suggesting that Unk was closely related or even identical to El-0 (Fig. 4B). Thereby, Unk and El-0 could be clearly distinguished from Hag-2 (ID9394) and Spro 3 (ID9452), the second best hits that agreed in only 107 genotypes (Suppl. Table 5). Again, this analysis was slightly biased due to missing genotypes. However, the informative value of such an analysis could be further increased, if required, by increasing the number of markers to the full 140SNPvCol marker set. Alternatively, a variant call format (vcf) file could be generated using the marker information provided in Suppl. Table 2 and the analysis could be run using the 1001 Genomes Strain ID tool (http://tools.1001genomes.org/strain_id/).

## Discussion

Genetic markers are important tools for genotyping and to study natural genetic variation. New methods implementing NGS technologies to obtain large scale genotypic information for analysis of mapping populations are on the rise (James et al., 2013; Rowan et al., 2015). Despite the availability of these attractive new methodologies that can provide the highest possible resolution, small-scale genotyping platforms are still required that provide solutions for flexible and cost-efficient genetic studies with minimal laboratory and bioinformatics effort. Here, we describe 140SNPvCol, a KASP-technology based SNP marker set, specifically designed to discriminate between the common reference accession Col-0 and the genotypes of most Arabidopsis accessions (Semagn et al., 2014). We successfully validated this marker set and integrated SNP data of the recently released 1,135 Arabidopsis genomic sequences (The 1001 Genomes Consortium, 2016). The 140SNPvCol marker set can be used for genetic mapping of phenotypic traits, marker-assisted backcrossing, the characterization of advanced backcross lines and genotyping mapping populations like recombinant inbred lines (RILs), specifically in cases where Col-0 was used as a parental genomic background. Further, it can be used to verify an accession’s identity or to assign its genetic group membership.

Many SNP genotyping platforms or SNP marker sets are not accessible to the broad community. Therefore, we decided to utilize the KASP genotyping platform as an economic and flexible alternative for mapping and genotyping in Arabidopsis. By developing and depositing the marker set at a service provider, we have made this marker set publicly available and users can take advantage of the provided genotyping service. Further, the 140SNPvCol assays can be custom-generated based on the provided sequence information (Suppl. Table 1).

Many genotyping platforms are not flexible in using individual subsets of marker sets and fixed experimental setups generate non-informative data that still involve costs. The 140SNPvCol KASP-based marker set, in turn, is flexible and cost efficient since subsets of the markers can be independently assembled and new markers can be developed and flexibly integrated into the 140SNPvCol set. In this way, the marker set can be optimally adapted to the genotyping needs of the specific experiment. We retrieved all 140SNPvCol genotypes of the 1,135 publically available genome sequences (The 1001 Genomes Consortium, 2016). The optimal 140SNPvCol subset of informative markers can be easily determined for each combination of Col-0 with any other accession or accessions using the information provided in Table S4. Therefore, the cost for genotyping is reduced to a minimum. In the same lines, we showed that ambiguously named 140SNPvCol genotypes of the 1001 Genomes SNP variant data can be determined by the 140SNPvCol markers. When calculated from the proportions of R and NR alleles, an overall high probability of around 87% is found that these alleles possess informative NR alleles. Therefore, we recommend to include the corresponding markers to the individual marker subset.

In conclusion, we generated a new resource for small-scale genotyping of SNP alleles in Arabidopsis. It is important to note, that the respective PCR assays to determine the marker genotypes were generated and are stored at a commercial service provider freely accessible to the community. Hence, cost is limited to genotyping per datapoint. As we demonstrated, the generation of this resource greatly benefited from the increasing availability of genomic sequence information. Based on the design of the marker set, we anticipate that also additional accessions of Arabidopsis sequenced in the future may be distinguished by these markers rendering them a unique resource with long term validity.

## Experimental procedures

### 1001 SNP data retrieval

SNP allele information of the 140SNPvCol marker set was retrieved from http://1001genomes.org/ and is provided in Suppl. Table 4.

### Marker information

Around 80 bp sequence, both upstream and downstream of each 140SNPvCol SNP is provided as Suppl. Table 1. The marker SNP allele is indicated as [R/NR] in line with the LGC Ltd. (Teddington, UK) SNP submission guidelines (https://www.lgcgroup.com/kasp/). Additional marker allele information is provided in Suppl. Table 2. All markers were submitted with the full name CK_SNP_[marker ID] to LGC Ltd. (Teddington, UK). Another LGC Ltd.-internal unique marker ID (LGC_SNPNum) is additionally provided in Suppl. Table 2.

### Pairwise distance calculations

The pairwise genetic distance was calculated with Geneious vR7.0.5 (Biomaters Limited, Auckland, New Zealand) using the HKY genetic distance model and the Neighbor-Joining tree build method. The number of identical alleles is reported; ambiguous SNP alleles were ignored. The input FASTA-format file is provided as Suppl. File 1. The reference R allele is deposited as A and the non-reference NR allele is deposited as B. The output pairwise genetic distance (as a number of shared alleles) is provided in Suppl. File 2.

### DNA extraction

DNA was extracted from a leaf sample using a standard phenol/chloroform extraction procedure. DNA was eluted in 100 μl nuclease-free water and DNA concentration was determined with a Nanodrop 2000 (ThermoFisher Scientific, Waltham, USA). All samples were equilibrated to a DNA concentration of 10 ng/μl with nuclease free water and 100 μl (required volume depends on the number of data points to be determined) were provided to LGC Ltd. (Teddington, UK). Further details and requirements of the LGC Limited genotyping service can be obtained from https://www.lgcgroup.com/kasp/

## Supplemental tables

**Supplemental Table 1:** 140SNPvCol marker sequences in the LGC^®^ SNP submission format.

**Supplemental Table 2:** Detailed information of the 140SNPvCol markers.

**Supplemental Table 3:** Genotypes of Col-0, Kil-0, L*er* analysed using the 140SNPvCol markers.

**Supplemental Table 4:** 140SNPvCol marker genotypes of 1,135 accessions, plain genotypes (A) and transformed to as R, NR, and N alleles (B)

**Supplemental Table 5:** 140SNPvCol genotype consensus between Unk and 1,135 accessions (A) and quantification thereof (B).

## Supplemental files

**Supplemental File 1:** Genotypes of 140SNPvCol SNPs of 1,135 accessions as FASTA file.

**Supplemental File 2:** Output of pairwise distance analysis.

## Acknowledgements

The authors would like to thank Thomas Regnault for discussions and critical reading of the manuscript and Paula Thompson for language editing. This work was supported by grants from the Deutsche Forschungsgemeinschaft as part of the SPP1530 “Flowering time control: from natural variation to crop improvement.” and the Sonderforschungsbereich 924 “Molecular mechanisms regulating yield and yield stability” to Claus Schwechheimer.

## References

Agrena, J., Oakley, C.G., McKay, J.K., Lovell, J.T., and Schemske, D.W. (2013). Genetic mapping of adaptation reveals fitness tradeoffs in Arabidopsis thaliana. Proc Natl Acad Sci U S A 110, 21077–21082.

Alonso-Blanco, C., Aarts, M.G., Bentsink, L., Keurentjes, J.J., Reymond, M., Vreugdenhil, D., and Koornneef, M. (2009). What has natural variation taught us about plant development, physiology, and adaptation? Plant Cell 21, 1877–1896.

The 1001 Genomes Consortium (2016). 1,135 genomes reveal the global pattern of polymorphism in *Arabidopsis thaliana*. Cell.

Gomaa, N.H., Montesinos-Navarro, A., Alonso-Blanco, C., and Pico, F.X. (2011). Temporal variation in genetic diversity and effective population size of Mediterranean and subalpine Arabidopsis thaliana populations. Mol Ecol 20, 3540–3554.

Gresham, D., Curry, B., Ward, A., Gordon, D.B., Brizuela, L., Kruglyak, L., and Botstein, D. (2010). Optimized detection of sequence variation in heterozygous genomes using DNA microarrays with isothermal-melting probes. Proc Natl Acad Sci U S A 107, 1482–1487.

James, G.V., Patel, V., Nordstrom, K.J., Klasen, J.R., Salome, P.A., Weigel, D., and Schneeberger, K. (2013). User guide for mapping-by-sequencing in Arabidopsis. Genome Biol 14, R61.

Konieczny, A., and Ausubel, F.M. (1993). A procedure for mapping Arabidopsis mutations using co-dominant ecotype-specific PCR-based markers. Plant J 4, 403–410.

Lukowitz, W., Gillmor, C.S., and Scheible, W.R. (2000). Positional cloning in Arabidopsis. Why it feels good to have a genome initiative working for you. Plant Physiol 123, 795- 805.

Lutz, U., Pose, D., Pfeifer, M., Gundlach, H., Hagmann, J., Wang, C., Weigel, D., Mayer, K.F., Schmid, M., and Schwechheimer, C. (2015). Modulation of ambient temperature-dependent flowering in *Arabidopsis thaliana* by natural variation of *FLOWERING LOCUS M*. PLoS Genet 11, e1005588.

Mendez-Vigo, B., Martinez-Zapater, J.M., and Alonso-Blanco, C. (2013). The flowering repressor SVP underlies a novel Arabidopsis thaliana QTL interacting with the genetic background. PLoS Genet 9, e1003289.

Pacurar, D.I., Pacurar, M.L., Street, N., Bussell, J.D., Pop, T.I., Gutierrez, L., and Bellini, C. (2012). A collection of INDEL markers for map-based cloning in seven Arabidopsis accessions. J Exp Bot 63, 2491–2501.

Peters, J.L., Cnudde, F., and Gerats, T. (2003). Forward genetics and map-based cloning approaches. Trends in plant science 8, 484–491.

Pico, F.X., Mendez-Vigo, B., Martinez-Zapater, J.M., and Alonso-Blanco, C. (2008). Natural genetic variation of Arabidopsis thaliana is geographically structured in the Iberian peninsula. Genetics 180, 1009–1021.

Platt, A., Horton, M., Huang, Y.S., Li, Y., Anastasio, A.E., Mulyati, N.W., Agren, J., Bossdorf, O., Byers, D., Donohue, K., Dunning, M., Holub, E.B., Hudson, A., Le Corre, V., Loudet, O., Roux, F., Warthmann, N., Weigel, D., Rivero, L., Scholl, R., Nordborg, M., Bergelson, J., and Borevitz, J.O. (2010). The scale of population structure in Arabidopsis thaliana. PLoS Genet 6, e1000843.

Rowan, B.A., Patel, V., Weigel, D., and Schneeberger, K. (2015). Rapid and inexpensive whole-genome genotyping-by-sequencing for crossover localization and fine-scale genetic mapping. G3 (Bethesda) 5, 385–398.

Schmid, K.J., Torjek, O., Meyer, R., Schmuths, H., Hoffmann, M.H., and Altmann, T. (2006). Evidence for a large-scale population structure of *Arabidopsis thaliana* from genome-wide single nucleotide polymorphism markers. Theor Appl Genet 112, 1104–1114.

Semagn, K., Babu, R., Hearne, S., and Olsen, M. (2014). Single nucleotide polymorphism genotyping using Kompetitive Allele Specific PCR (KASP): overview of the technology and its application in crop improvement. Molecular Breeding 33, 1–14.

Simon, M., Simon, A., Martins, F., Botran, L., Tisne, S., Granier, F., Loudet, O., and Camilleri, C. (2012). DNA fingerprinting and new tools for fine-scale discrimination of Arabidopsis thaliana accessions. Plant J 69, 1094–1101.

Simon, M., Loudet, O., Durand, S., Berard, A., Brunel, D., Sennesal, F.X., Durand-Tardif, M., Pelletier, G., and Camilleri, C. (2008). Quantitative trait loci mapping in five new large recombinant inbred line populations of Arabidopsis thaliana genotyped with consensus single-nucleotide polymorphism markers. Genetics 178, 2253–2264.

Tabone, T., Mather, D.E., and Hayden, M.J. (2009). Temperature switch PCR (TSP): Robust assay design for reliable amplification and genotyping of SNPs. BMC Genomics 10, 580.

Thomson, M. (2014). High-Throughput SNP Genotyping to Accelerate Crop Improvement. Plant Breeding and Biotechnology, 195–212.

Torjek, O., Berger, D., Meyer, R.C., Mussig, C., Schmid, K.J., Rosleff Sorensen, T., Weisshaar, B., Mitchell-Olds, T., and Altmann, T. (2003). Establishment of a high-efficiency SNP-based framework marker set for Arabidopsis. Plant J 36, 122–140.

Weigel, D. (2012). Natural variation in Arabidopsis: from molecular genetics to ecological genomics. Plant Physiol 158, 2–22.

Weigel, D., and Nordborg, M. (2005). Natural variation in Arabidopsis. How do we find the causal genes? Plant Physiol 138, 567–568.

Wijnker, E., van Dun, K., de Snoo, C.B., Lelivelt, C.L., Keurentjes, J.J., Naharudin, N.S., Ravi, M., Chan, S.W., de Jong, H., and Dirks, R. (2012). Reverse breeding in Arabidopsis thaliana generates homozygous parental lines from a heterozygous plant. Nat Genet 44, 467–470.

Yuan, W., Flowers, J.M., Sahraie, D.J., Ehrenreich, I.M., and Purugganan, M.D. (2016). Extreme QTL mapping of germination speed in Arabidopsis thaliana. Mol Ecol 25, 4177–4196.

